# Environment of origin and domestication affect morphological, physiological, and agronomic response to water deficit in chile pepper (*Capsicum* sp.)

**DOI:** 10.1101/2021.11.16.468826

**Authors:** J.E. McCoy, L. McHale, M. Kantar, L. Jardón-Barbolla, K.L. Mercer

## Abstract

Global climate change is having a significant effect on agriculture by causing greater precipitation variability and an increased risk of drought. To mitigate these effects, it is important to identify specific traits, adaptations, and germplasm that improve tolerance to soil water deficit. Local varieties, known as landraces, have undergone generations of farmer-mediated selection and can serve as sources of variation, specifically for tolerance to abiotic stress. Landraces can possess local adaptations, where accessions adapted to a particular environment will outperform others grown under the same conditions. We explore adaptations to water deficit in chile pepper landraces from across an environmental gradient in Mexico, a center of crop domestication and diversity, as well in improved varieties bred for the US. In the present study, we evaluated 25 US and Mexico accessions in a greenhouse experiment under well-watered and water deficit conditions and measured morphological, physiological, and agronomic traits. Accession and irrigation regime influenced biomass and height, while branching, CO2 assimilation, and fruit weight were all influenced by an interaction between accession and irrigation. *A priori* group contrasts revealed possible adaptations to water deficit for primary branching, CO_2_ assimilation, and plant height associated with geographic origin, domestication level, and pepper species. Additionally, within the Mexican landraces, the number of primary branches had a strong relationship with precipitation from the environment of origin. This work provides insight into chile pepper response to water deficit and adaptation to drought and identifies possibly tolerant germplasm.

## Introduction

Global temperatures are rising due to anthropogenic climate change, causing greater precipitation variability and an accelerated risk of drought [1]. Increasing incidence of drought has considerable implications for agriculture. Crop genetic improvement, such as breeding of varieties capable of tolerating drought, could mitigate the effects of climate change on agriculture [2]. In order to do this, we must first identify sources of diversity and specific traits associated with drought adaptation and water deficit tolerance. Landraces, i.e., local crop varieties from centers of diversity, have undergone less intentional breeding, thus representing a different level of domestication than commercial cultivars and may serve as sources for biotic and abiotic stress tolerance [3–5]. Yet, there is also growing evidence that substantial diversity can be maintained under conventional breeding, as seen recently in tomato [6]. Still, landraces likely possess higher variation for response to abiotic stress and require further exploration, alongside their more commercial counterparts. Additionally, this understanding of variation between commercial and landrace germplasm could converge to inform participatory plant breeding programs aimed at addressing local farmers needs for climate change adaptation [7,8].

Chile pepper (*Capsicum* sp.) is an economically and culturally significant crop worldwide, with over 36 million tons harvested in 2017 [9]. Mexico is the second largest producer and the center of domestication and diversity for chile pepper [9–11]. Near the center of domestication and diversity, crops and their wild relatives may possess adaptations to local conditions [12]. These adaptations may result from genetic differentiation for traits affecting productivity, such as the phenotypic responses to varied growing environments [13]. For example, in their local environments, Mexican maize (*Zea mays* L. subsp. *mays*) landraces from a given elevation tend to demonstrate higher overall fitness than non-local ones; local landraces also tend to outperform improved varieties under local conditions [14]. In wild and landrace chile pepper, local adaptation may also be possible, where individuals (referred to as accessions) originating from low precipitation environments may possess elevated tolerance to soil water deficit. For example, a recent germination study identified adaptive responses in chile pepper associated with precipitation [15]. Landraces from hotter, drier ecozones had significantly delayed germination, suggesting a possible adaptive response related to drought avoidance. Evidence of significant genetic diversity exists in modern pepper germplasm in the US, particularly for biotic stress tolerance[e.g. 16,17]; however, landrace germplasm likely possesses more genetic variation. For this reason, it is important to explore tolerance to soil water deficit in both US and Mexican accessions.

This greenhouse study evaluated the response to soil water deficit in a diverse set of germplasm selected from US and Mexican origins. The objectives of this experiment were to: (1) measure the morphological, physiological, and agronomic response of diverse chile pepper to soil water deficit; (2) evaluate differences between accessions, including differences associated with environmental and geographic origin and domestication gradients; and (3) identify unique responses of accessions to soil water deficit (i.e., genotype by environment interactions) that may indicate a tolerance to soil water deficit. Results of this study offer insight into adaptations to drought in chile pepper and highlight germplasm for continued research in water deficit tolerance.

## Materials and Methods

### Plant material

Our 25 chile pepper accessions included 18 from the US and seven from Mexico (Table 1). U.S. accessions were no longer under USDA plant variety protection status and had diverse plant habit, fruit type, and origin (Table 1). Specific US accessions included one commercial bell, one cherry, one *chilhuacle* landrace from Mexico that is produced in and selected for the Northwestern US, three commercial *jalapeño* cultivars, six New Mexican cultivars or landraces, one ornamental, two paprika (one US bred and one bred in Hungary, both produced in the US), one Mexican *serrano* type that is produced in the US, and two sweet peppers.

**TABLE 1.** List of germplasm used in a greenhouse soil water deficit experiment on chile pepper at the Ohio State University. Accessions are organized by country of origin.

| Name<sup>a</sup> | Species<sup>b</sup> | Pod Type | Seed Source | Origin Description |
| --- | --- | --- | --- | --- |
| U.S. Germplasm |  |  |  |  |
| Kaala | *C. annuum* | Bell | UHM Seed Lab | Developed and produced by University of Hawaii |
| PI 592813 | *C. annuum* | Cherry | NPGS | Maintained by the plant Genetic Resources Conservation Unit, Georgia |
| Chilhuacle Negro | *C. annuum* | Chilhuacle | Adaptive Seeds | Mexican landrace, produced in Oregon |
| Waialua | *C. annuum* | Jalapeño | UHM Seed Lab | Developed and produced by University of Hawaii |
| Tam Jalapeño | *C. annuum* | Jalapeño | Reimer Seeds | Developed by Texas A&M University, produced in Maryland |
| Tam Vera Cruz | *C. annuum* | Jalapeño | Reimer Seeds | Developed by Texas A&M University, produced in Maryland |
| Canoncito | *C. annuum* | New Mexican | Wild Garden Seed | New Mexican landrace, produced in Oregon |
| Chimayo | *C. annuum* | New Mexican | Adaptive Seeds | New Mexican landrace, produced in Oregon |
| Anaheim M | *C. annuum* | New Mexican | Reimer Seeds | U.S. cultivar, produced in Maryland |
| Anaheim TMR 23 | *C. annuum* | New Mexican | Reimer Seeds | U.S. cultivar, produced in Maryland |
| PI 586666 | *C. annuum* | New Mexican | NPGS | Maintained by the plant Genetic Resources Conservation Unit, Georgia |
| PI 159229 | *C. annuum* | New Mexican | NPGS | Maintained by the plant Genetic Resources Conservation Unit, Georgia |
| PI 631153 | *C. annuum* | Ornamental | NPGS | Maintained by the plant Genetic Resources Conservation Unit, Georgia |
| Szegedi 179 | *C. annuum* | Paprika | Adaptive Seeds | Originated in Hungary, produced in Oregon |
| NuMex Conquistador | *C. annuum* | Paprika | Reimer Seeds | Developed by New Mexico State University, produced in Maryland |
| Hidalgo Hot | *C. annuum* | Serrano | Reimer Seeds | Originated in Mexico, produced in Maryland |
| Stocky Golden Roaster | *C. annuum* | Sweet | Wild Garden Seed | Farm-original variety, Oregon |
| PI 586665 | *C. annuum* | Sweet | NPGS | Maintained by the plant Genetic Resources Conservation Unit, Georgia |
| Mexico Germplasm |  |  |  |  |
| Ca0057 | *C. annuum* | Chile de Agua | OSU Collection | Collected from Oaxaca central valley plantation |
| Ca0256 | *C. annuum* | Chilgole | OSU Collection | Collected from Oaxaca coast backyard |
| Ca0045 | *C. annuum* | Costeño Rojo | OSU Collection | Collected from Oaxaca coast plantation |
| Ca0310 | *C. annuum* | Tusta | OSU Collection | Collected from Oaxaca central valley Letstand |
| Ca0344 | *C. annuum* | Tusta | OSU Collection | Collected from Oaxaca central valley letstand |
| Cc0144 | *C. chinense* | Habanero | OSU Collection | Collected from Oaxaca coast backyard |
| Cf0173 | *C. frutescens* | Mirasol | OSU Collection | Collected from Oaxaca coast backyard |

<sup>a</sup>Names refers to given cultivar, landrace name, or database accession number used for USDA and Mexico germplasm.
<sup>b</sup>Genus for all species is *Capsicum*.

We chose the seven Mexican accessions based on results from an environmental association analysis conducted on chile peppers collected from diverse environments in the Mexican state of Oaxaca [18]. Using environment of origin as a “phenotype” in an environmental association analysis [see 19], Bernau [18] identified single nucleotide polymorphisms (SNPs) associated with specific environments of origin. Using the identified SNPs, we selected seven Mexican accessions with alleles associated with drought-prone environments; some had also shown themselves to have some tolerance to water deficit in the greenhouse. Mexican accessions included five *C. annuum* – one *chile de agua* from a central valley plantation, one *chilgole* from a costal backyard, one *costeño rojo* from a costal plantation, and two *tusta* semi-wild from the central valley – and two additional species. The *C. chinense* and *C. frutescens* accessions were a *habanero* and *mirasol*, respectively, and had been collected from costal backyards.

### Study system and experimental design

We grew the 25 chile pepper accessions under greenhouse conditions at the Ohio State University in Columbus, Ohio. Seedlings were transplanted into 6 L pots using BM-1 Potting Mix (Berger, Saint-Modeste, QC, Canada) approximately six weeks after planting. Slow release, 14-14-14 fertilizer (Osmocote, Scotts Miracle-Gro, Marysville, OH, USA) was mixed into the soil at transplanting and three fertilizer applications of a 20-10-20 Peat-Lite fertilizer (JR Peters Inc., Allentown, PA, USA) were applied through irrigation lines, as needed.

We implemented a randomized complete block design with four blocks; individual greenhouse bench serving as our blocking factor. Within each block we applied factorial combinations of accession (25 levels, see above) and irrigation (water deficit and control, see below) to each pot (one plant per pot) or experimental unit.

Plants were subjected to two levels of irrigation: daily watering (control) and weekly watering (water deficit). Water application varied slightly throughout the growing season to accommodate light, temperature, and transpiration rate changes; however, plants were watered to saturation at each watering event. Stomatal conductance and CO_2_ assimilation were measured weekly using the LI-6800 Portable Photosynthesis System (Licor, Lincoln, NE, USA). Instrument conditions were as follows: Flow rate 600 µmol/s, relative humidity 65%, CO_2__s 400 ppm, fan speed 10,000 rpm, and fluorometer adjusted to ambient light conditions. Prior to logging a measurement, the LI-6800 was stabilized on relative humidity, stomatal conductance, and CO_2_ assimilation. After approximately four months, plants were destructively harvested and measurements of morphological and agronomic traits were collected, including: plant height, above-ground biomass, number of primary branches, fruit weight, and 100-seed weight. Due to the varying phenology within Mexican accessions, many accessions were still fruiting as the experiment was terminated. Thus, we only report on analyses of fruit traits for US accessions.

### Data analysis

To perform three different sets of analyses, we used the lmerTest package in R version 4.0.5 [20]. First, to assess the effect of experimental manipulations on our response variables, we employed a general linear mixed model for analysis of variance (ANOVA). For growth, morphological, and physiological metrics, we analyzed data in all accessions with irrigation, accession, and their interaction as fixed effects, and block as a random effect. Due to limitations on our fruit data in Mexican accessions (see above), we analyzed just the 18 U.S. cultivars to assess agronomic performance (fruit weight, seed weight). For all analyses, ANOVA assumptions were checked. Log transformations were performed on CO_2_ assimilation and stomatal conductance due the heteroscedastic nature of the data. For significant effects, we generated mean separation tables using Tukey’s HSD test.

Second, to discern effects of country of origin, landrace status, and *Capsicum* species, we conducted a series of *a priori* contrasts within the previous model. To measure the effect of country of origin, we contrasted Mexican and U.S. germplasm. We opted to contrast landrace status because several U.S. accessions in this experiment are marketed as either U.S. or Mexican landraces. Though not collected directly from Mexico, it is possible that these accessions may have undergone less commercial breeding and subsequently possess adaptations to abiotic stress [5]. Thus, we contrasted landraces (both US and Mexico) with others. Since plant habit can differ greatly across species, we contrasted our *C. chinense* (Cc0144) and our *C. fructescens* (Cf0173) accessions with our *C. annuum* ones (Table 1) to better understand differences due to speciation.

Third, to address the relationship between environment of origin and response to water deficit in our collected traits, we performed regression analysis using only Mexican accessions. Our collection of Mexican accessions are georeferenced and thus can be analyzed for specific environmental variables from their originating location using the publicly available Bioclim data and soil data from the ISRIC [21,22]. We selected variables directly related to precipitation, including total available soil water capacity, BIO12 (annual mean precipitation), BIO15 (precipitation seasonality), BIO16 (precipitation of the wettest quarter), BIO17 (precipitation of the driest quarter), BIO18 (precipitation of the warmest quarter), and BIO19 (precipitation of the coldest quarter). In order to reduce the overall size of the linear model, we generated a correlation matrix of the seven environmental variables (Fig S1). Using the corresponding P values and correlation coefficients (data not shown), we identified all significant correlations (α = 0.05, coefficient > 0.80) and determined that a linear model using total available soil water, annual mean precipitation, and precipitation seasonality would be representative of the full model. The experimental blocking factor contributed very little to variation in the model and was confirmed to be unnecessary in all models based on lower Akaike information criterion (AIC) scores for models without block. Ultimately, we used a linear regression model including the three environmental variables (total available soil water, annual mean precipitation, and precipitation seasonality) plus irrigation treatment and its interaction with each variable.

## Results

### Response to water deficit

Plant biomass and plant height were significantly affected by water treatment and were reduced by an average of 10.6 g (16.3%) and 7.6 cm (12.3%), respectively, under water deficit (Tables 2 & S1). Mean Plant biomass and height were also significantly affected by accession (Tables 2 & 3). Plant biomass means ranged significantly among accessions from 49.6 g (SE = 3.30; Szegedi 179) to 75.7 g (SE = 3.30; Ca0344); plant height ranged from 36.2 cm (SE = 4.30; PI631153) to 81.2 cm (SE = 4.30; *Chilhuacle Negro*) (Table 3).

**TABLE 2.**
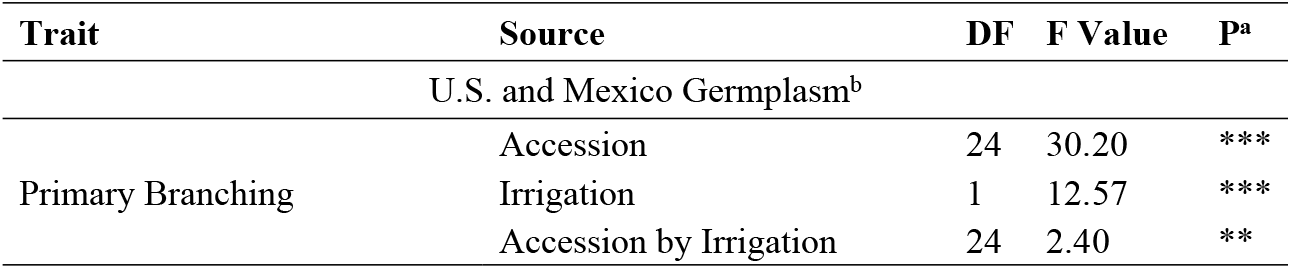
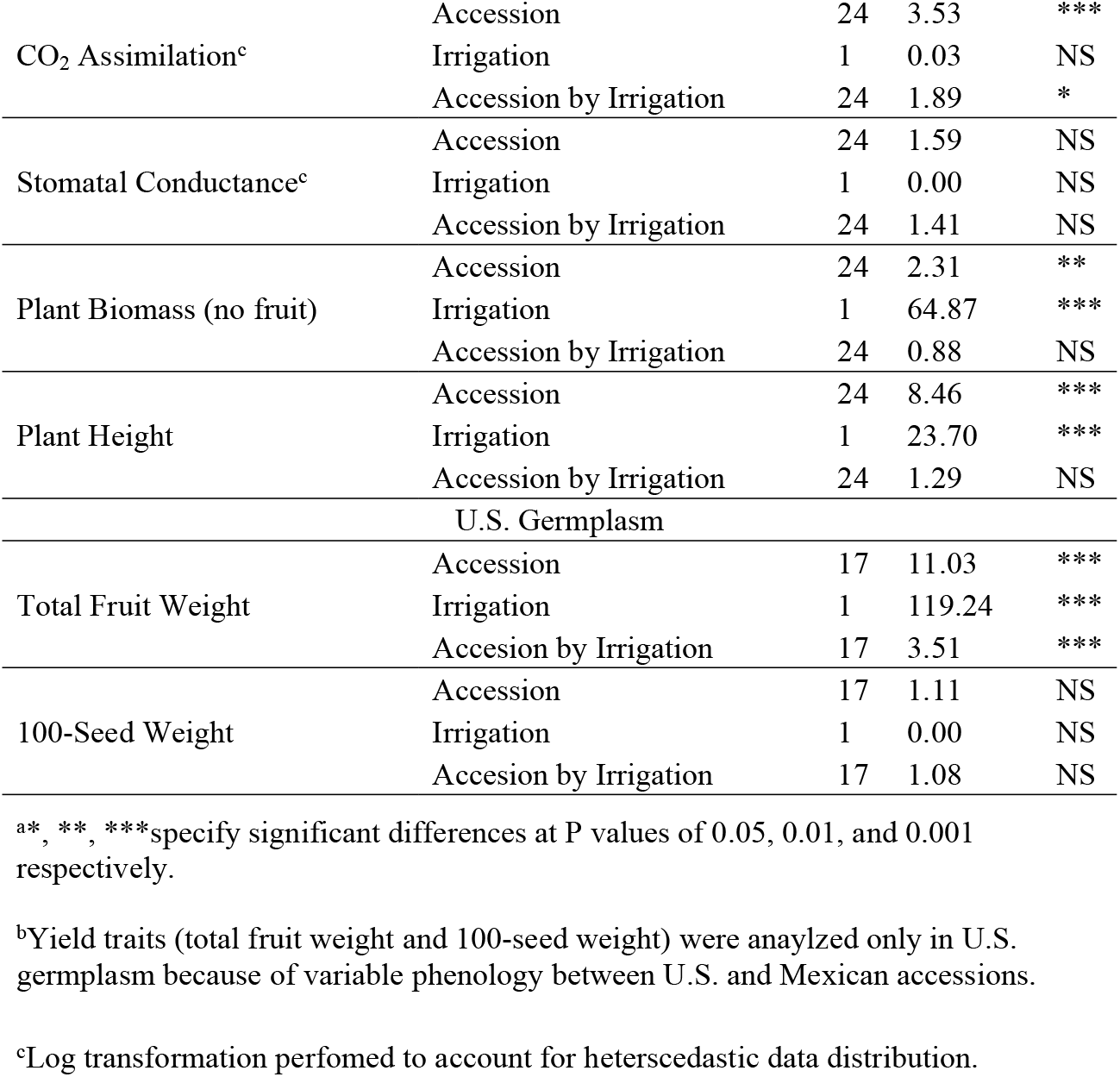
Analysis of variance for seven traits in a greenhouse soil water deficit experiment on chile pepper (*Capsicum* sp.) at the Ohio State University. Data was analyzed in R (version 4.0.5) using a linear mixed model. Source was considered significant if P < 0.05.

**TABLE 3.**
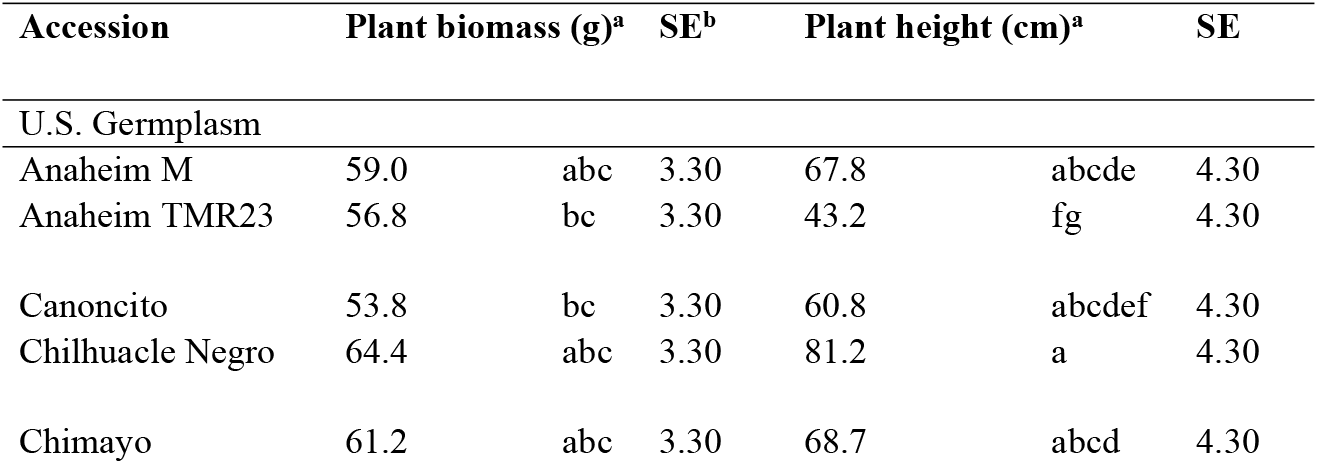
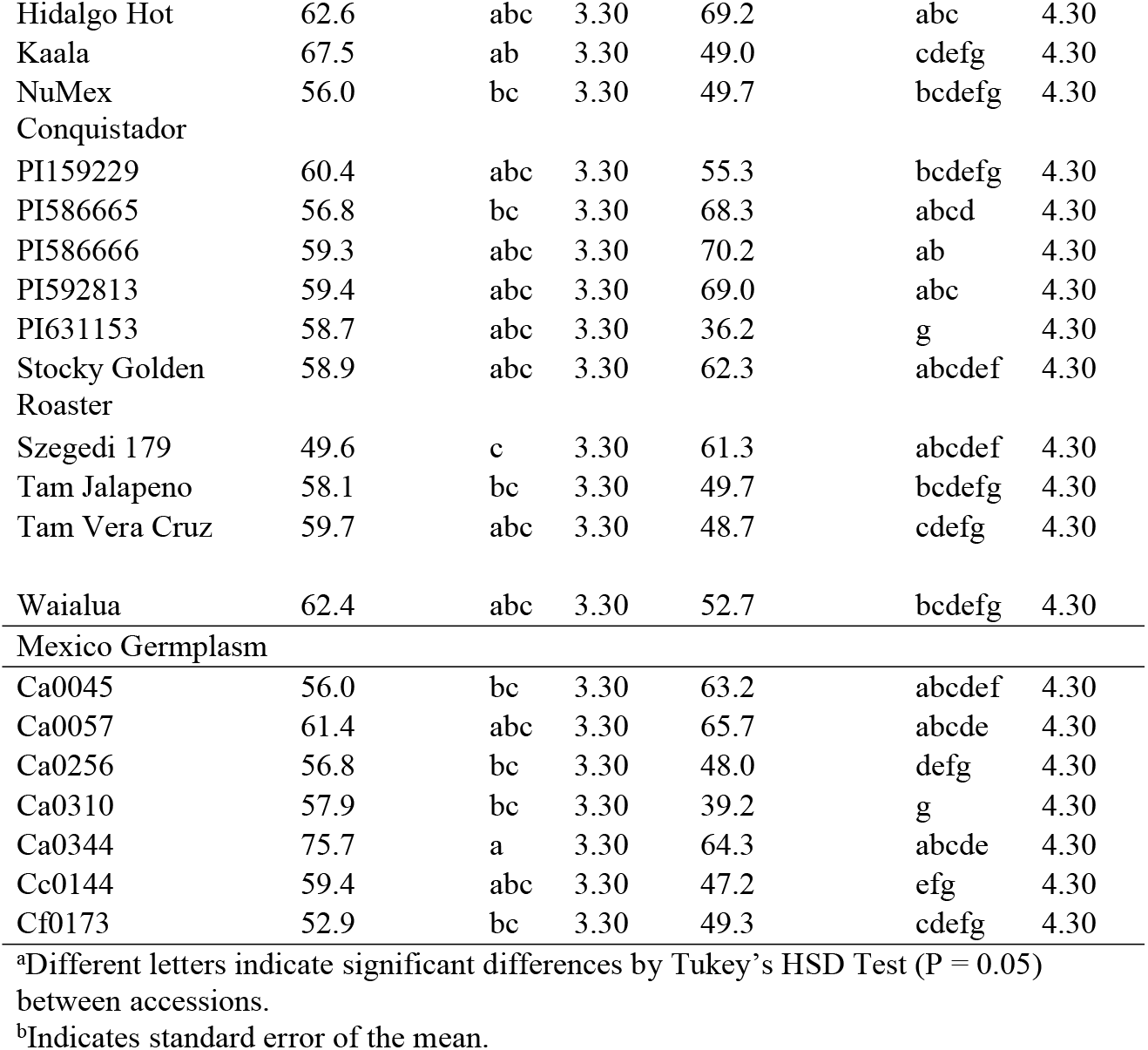
Mean separation (estimated marginal means) of two traits with significant main affects for accession from a greenhouse soil water deficit experiment on chile pepper (*Capsicum* sp.) at the Ohio State University. Organized by country of origin.

A significant interaction between accession and irrigation for primary branching and CO_2_ assimilation indicated a differential response to water deficit among accessions (Table 2). Many accessions experienced decreased branching under water deficit, but to differing degrees (Fig 1). Surprisingly, the USDA accession PI631153 had a great increase in number of primary branches under water deficit. CO_2_ assimilation rates in commercial US accessions either declined, stayed the same, or increased under water deficit (Fig 2). However, CO_2_ assimilation rates in landrace accessions (from US and Mexico) either remained the same or increased under water deficit, but never declined.

**FIGURE 1.**
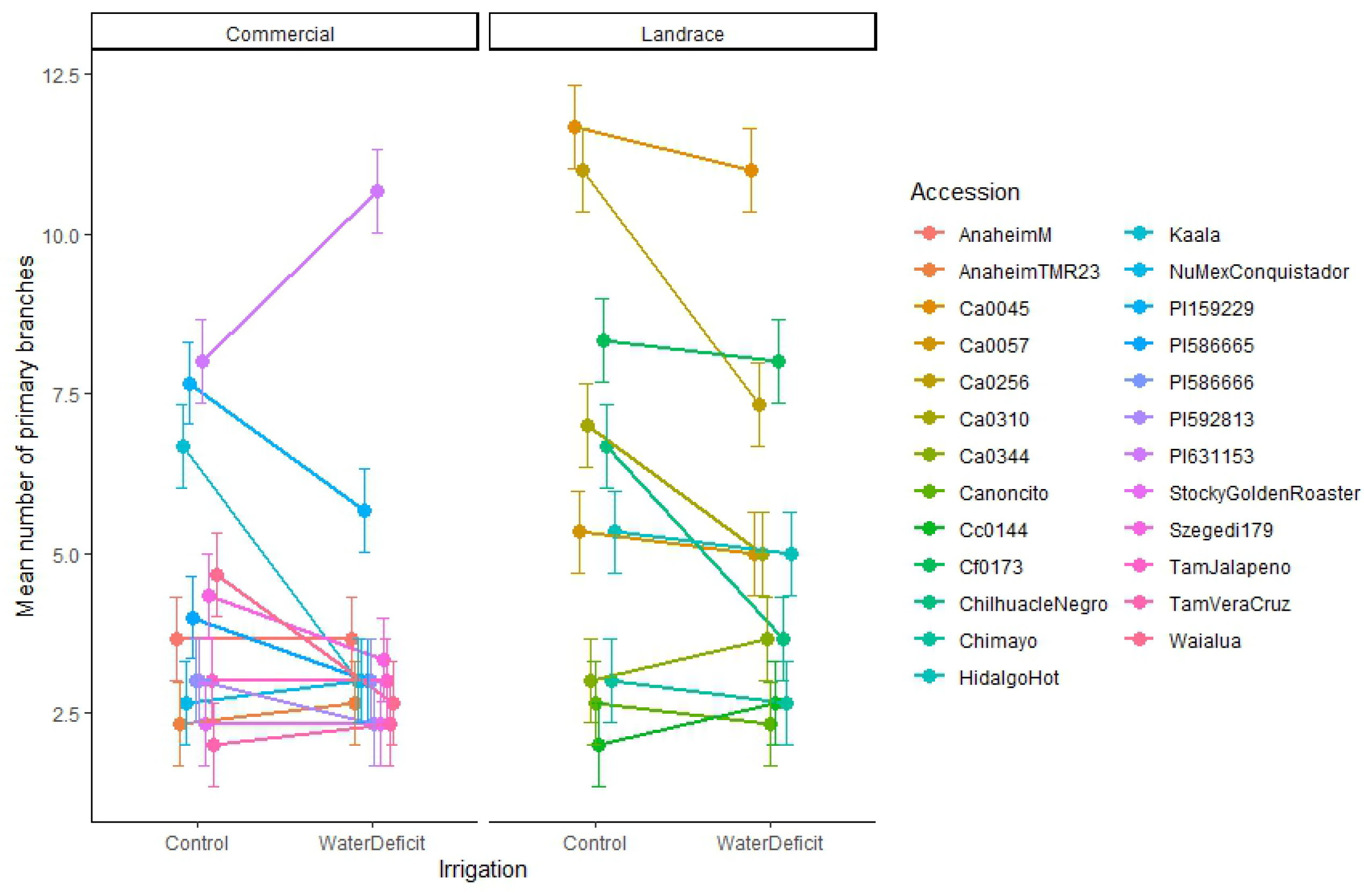
Interaction plot of estimated marginal means with standard error bars for primary branch number in a greenhouse soil water deficit experiment on chile pepper (*Capsicum* sp.) at the Ohio State University. Accessions are separated by domestication.

**FIGURE 2.**
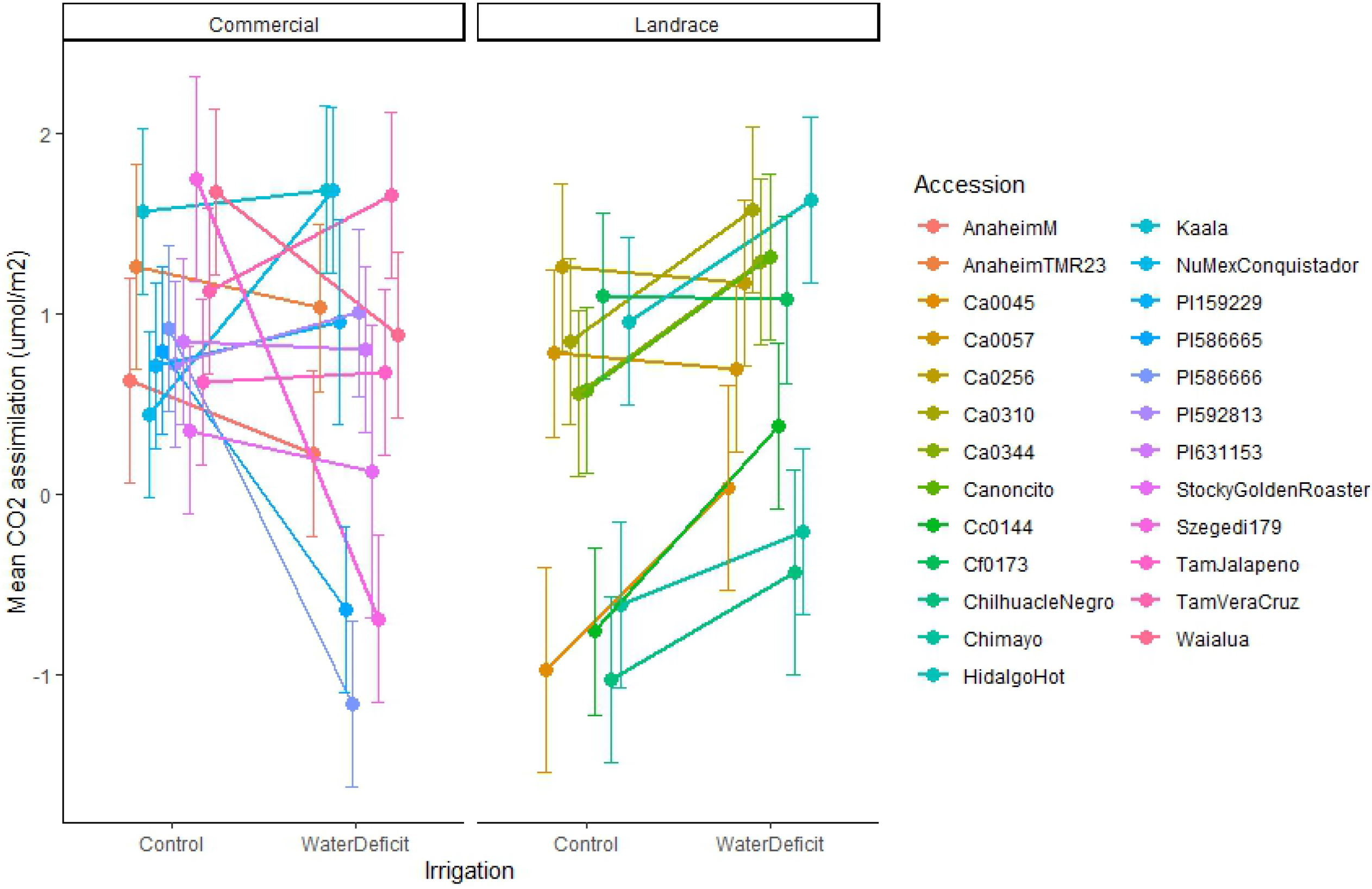
Interaction plot of estimated marginal means with standard error bars for CO_2_ assimilation (log transformed) in a greenhouse soil water deficit experiment on chile pepper (*Capsicum* sp.) at the Ohio State University. Accessions are separated by domestication.

Focusing only on US accessions for agronomic traits, we found a significant variation in yield response to water deficit among accessions (Table 2; Fig 3). In general, mean total fruit weight was reduced under water deficit, but to varying degrees. Anaheim M had the greatest total fruit yield under well-watered conditions but had a substantial decrease under water deficit. Alternatively, *Canoncito* had no significant difference in total fruit weight between levels of irrigation, suggesting possible tolerance (Fig 3).

**FIGURE 3.**
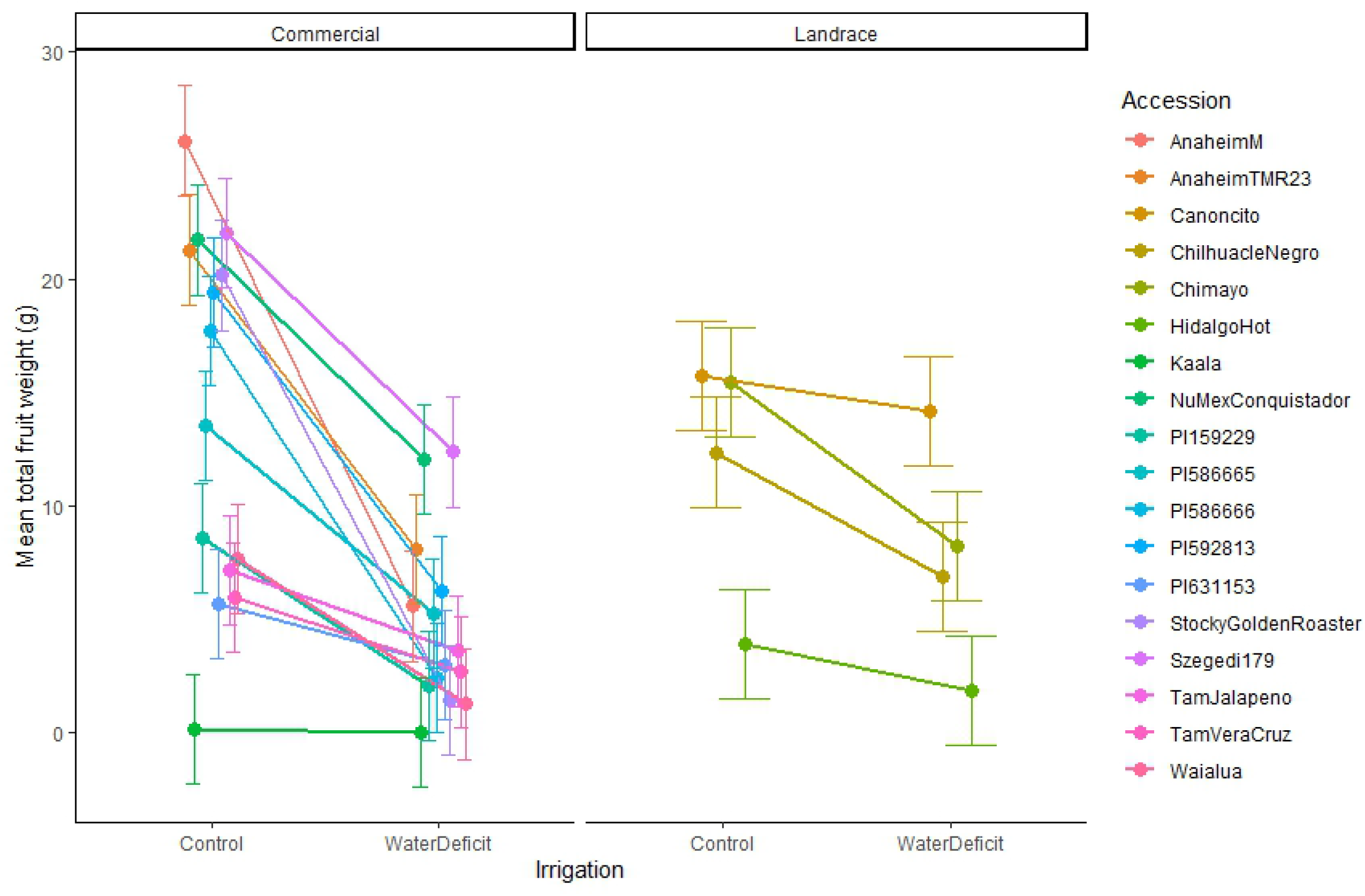
Interaction plot of estimated marginal means with standard error bars for mean fruit weight in U.S. accessions only in a greenhouse soil water deficit experiment on chile pepper (*Capsicum* sp.) at the Ohio State University. Accessions are separated by domestication.

### Environment of origin and domestication effects

Examining our *a priori* contrasts, we found significant differences between groups Mexico vs. US, Landrace vs. other, and Annuum vs. other (Table 4). Under both well-watered and water deficit conditions, primary branching was significantly higher in Mexican accessions than US accessions and in landraces as compared to improved varieties (Table 4). Primary branching was significantly lower in *C. annuum* than in other species under water deficit, but not under well-watered conditions (Table 4). Plants from the US were taller than those from Mexico under well-watered conditions, but we found no differences between the two under water deficit (Table 4). Additionally, *C. annuum* accessions were taller than accessions from other species under well-watered conditions, but that difference declined under water deficit (Table 4).

**TABLE 4.** Results of *a priori* group contrasts on four traits from a greenhouse soil water deficit experiment on chile pepper (*Capsicum* sp.) at the Ohio State University.

| Trait | Mexico:US<sup>a</sup> |  |  |  | Landrace:Other |  |  |  | Annuum:Other |  |  |  |
| --- | --- | --- | --- | --- | --- | --- | --- | --- | --- | --- | --- | --- |
|  | Control |  | Water Deficit |  | Control |  | Water Deficit |  | Control |  | Water Deficit |  |
|  | t value | P<sup>b</sup> | t value | P | t value | P | t value | P | t value | P | t value | P |
| Primary Branching | 9.44 | ***<br>> | 8.69 | ***<br>> | 7.26 | ***<br>> | 5.72 | ***<br>> | -0.53 | NS | -2.38 | * < |
| CO<sub>2</sub> Assimilation | -1.58 | NS | 1.40 | NS | -3.69 | ***<br>< | 0.95 | NS | 1.52 | NS | -0.18 | NS |
| Plant Biomass | 1.62 | NS | -0.77 | NS | 1.88 | NS | -0.34 | NS | 0.24 | NS | 1.78 | NS |
| Plant Height | -2.38 | * < | -1.88 | NS | 1.87 | NS | 1.5 | NS | 3.05 | ** > | 1.95 | NS |

<sup>a</sup>Contrasts were made in the following three groups: Mexico vs. U.S. accessions, Landrace accessions vs. others, and *C. annuum* vs. others.
<sup>b</sup>\*, \*\*, \*\*\*specify significant differences at P values of 0.05, 0.01, and 0.001 respectively. Greater than/less than symbols indicate the direction of the contrast vis-à-vis the order of the groups in the heading.

CO_2_ assimilation was significantly lower in landraces than in other accessions when well-watered. Yet, we found no differences under water deficit (Table 4). This reinforces the previously mentioned significant interactions between accession and irrigation on CO_2_ assimilation, indicating that assimilation rates were maintained within landrace accessions across irrigation treatments, but not in US accessions.

Lastly, results of the multiple regression analysis suggest that key environmental parameters related to precipitation are significant predictors of branching (Table 5). More specifically, we found a significant negative linear relationship between branching and total available soil water and precipitation seasonality (Table 5). Each unit increase in soil water content predicted a decrease in 2.699 branches (SE = 0.430, P ≤ 0.001) and each unit increase in precipitation seasonality predicted a decrease in 0.214 branches (SE = 0.073, P = 0.006). We identified a significantly positive relationship between annual mean precipitation and primary branching, where each unit of increase in precipitation predicts an increase in 0.014 branches (SE = 0.002, P ≤ 0.001). Contrary to results of the ANOVA, irrigation treatment did not have a significant relationship with primary branching. This suggests that environment plays a more significant role than water regime in predicting primary branching. Overall, the model accounted for 69.7% of the variation in the data (Table 5).

**TABLE 5.** Multivariate regression analysis of Mexican chile pepper accessions from a greenhouse soil water deficit experiment on chile pepper (*Capsicum* sp.) at the Ohio State University. The relationship between mean primary branching and environmental parameters related to precipitation from the originating environment are presented (R^2^ = 0.697).

| Trait | Predictors | Beta<sup>a</sup> | SE<sup>b</sup> | P<sup>c</sup> |
| --- | --- | --- | --- | --- |
| Primary<br>Branching | Irrigation | -14.803 | 33.451 | NS |
|  | Total Available Soil Water | -2.699 | 0.430 | *** |
|  | Annual Mean Precipitation | 0.014 | 0.002 | *** |
|  | Precipitation Seasonality | -0.214 | 0.073 | ** |
|  | Irrigation by Total Available Soil Water | 0.328 | 0.609 | NS |
|  | Irrigation by Annual Mean Precipitation | -0.003 | 0.003 | NS |
|  | Irrigation by Precipitation Seasonality | 0.023 | 0.103 | NS |

<sup>a</sup>Beta estimate indicates the change in primary branching relative to one unit change of the predictor.
<sup>b</sup>Indicates standard error of the mean.
<sup>c</sup>\*, \*\*, \*\*\*specify significant relationship at P values of 0.05, 0.01, and 0.001 respectively.

## Discussion

We found clear variation in responses of chile pepper accessions to soil water deficit, confirming that variation in tolerance may exist in germplasm currently in use. Much of the plant response to water deficit pointed to clear reductions in growth, but primary branching and CO_2_ assimilation did not respond consistently to water deficit across accessions. In fact, Mexican accessions, and landraces overall, better maintained CO_2_ assimilation under water deficit than their improved counterparts. Moreover, Mexican accessions branched more than US accessions, landraces were branchier than improved accessions, and *C. annuum* branched less than other species when well-watered. Finally, we found precipitation of the originating environment to predict branching in Mexican landraces. Thus, variation in physiological and growth form attributes may point to adaptations in pepper.

### Influence of water deficit on plant architecture and physiology

The wide range of primary branching within our accessions and the varying response of primary branching to water deficit suggests that plant architecture plays an important role in plant adaptation to drought. Branching may have biological significance relative to response to soil water deficit, through canopy management for optimal transpiration rates and shoot to root biomass partitioning, for example [see 23,24]. However, branching likely plays a specific role in water deficit response that goes beyond biomass partitioning, as above-ground biomass did not follow the same trends (i.e. no interaction between irrigation and accession). Leaf area index (LAI), the ratio of leaf area to ground area, plays a significant role in plant photosynthesis and generally decreases under drought [23]. Drought stress not only reduces leaf area, but also the photosynthetic rate per unit leaf area [25]. Shifts in plant architecture through additional branching may optimize overall leaf area, therefore, counteracting some of the reduced area caused by water stress. In addition to leaf area management, increased branching may help reduce photosynthetic stress brought on by reduced productivity associated with drought. Under drought stress, photosynthetic reactions are reduced and there is an the accumulation of damaging reactive oxygen species (ROS) [25]. There are several metabolic adaptations for reducing this damage, termed photoprotection, that help the plant dissipate excessive thermal energy [26]. Though we did not test for metabolic adaptations, it is possible that greater branching could provide a physical barrier, increasing diffuse light and decreasing direct light penetrating the canopy, subsequently reducing the stress of excessive photosynthetic energy under stress. Branching undoubtedly influences adaptation to water deficit, but without additional data on leaf area, it is difficult to fully explain its role here.

In chile pepper, the influence of water deficit on branching is variable and dependent on the germplasm. Showemimo and Olarewaju [27] found that branching in an advanced pepper breeding line decreased with severity of drought applied whereas we saw inconsistent responses across our diverse germplasm. However, the advanced breeding line response is similar to that observed in improved, US accessions overall (Table 4, Landrace vs. Other). Bernau [18], using landraces from southern Mexico, indicated that branching may relate to environmental adaptation in this system. Branching was lower in chile pepper accessions from the wet, western coast of Oaxaca and higher in accessions from the drier ecozones, Eastern Oaxaca and the Yucatan peninsula. Interestingly, Bernau also found branching decreased in accessions from more intensively cultivated systems (i.e. was higher in forest accessions and lower in plantation accessions). Morphological plasticity in response to water deficit has been shown to differ across levels of domestication. In seven species including *Beta vulgaris, Brassica oleracea, Helianthus annuus, Solanum lycopersicum, Triticum durum, Zea mays*, and *Pisum sativum*, norms of reaction differed in response to water deficit, where domesticated accessions greatly decreased performance, but their wild progenitors did not have as severe a response [28]. While this could explain the differing response between US and Mexican accessions, further investigation on branching across a domestication gradient is required.

Similar to branching, the differential CO_2_ assimilation responses we found across accessions indicate possible physiological adaptations to drought. The general expectation of plants under water stress is that assimilation rates decrease; however, the relationships between CO_2_ assimilation, stomatal conductance, and overall photosynthetic activity are tightly connected and complex [29,30]. Under mild water stress, net assimilation rates decrease as a result of decreased stomatal conductance. It is important to acknowledge that the gas exchange measurements we collected represented the activity in a single leaf at a given point in time. This snap-shot provides valuable insight into plant response to water deficit, but it does not explain whole-plant dynamics. Leaf area and distribution, in addition to CO_2_ assimilation per unit area, is a significant detail for understanding plant growth under stress [30]. In fact, the interactions found for primary branching and CO_2_ assimilation may be closely linked, where changes in architecture influenced whole-plant gas exchange. As previously mentioned, branching may alter light capture in the canopy, with increased number of branches having the potential to decrease stress caused by excessive light and thermal energy. Alternatively, changes to leaf area as a result of branching will influence photosynthetic activity and water use efficiency (WUE). WUE differs among genotypes, but it is complex at the leaf level [31]. However, Hatfield and Dole [31] highlight that differences in WUE among genotypes are related to net CO_2_ assimilation, leaf area, and specific leaf area. Another morphological change affecting physiological response to water deficit that is difficult to account for is stomatal density and size. Natural variation in stomatal density and size exists and has a direct influence on WUE, as seen in Arabidopsis [32].

### Influence of environment of origin and domestication on plant response to water deficit

We observed a geographic (i.e. Mexico vs. US) and domestication (i.e. landrace vs. other) effect on plant architecture, where primary branching was highest in Mexican accessions and landraces (both from Mexico and the US) across irrigation treatments (Table 4). Interestingly, we found that some accessions, particularly landraces, either maintained or even increased assimilation rates under water deficit (Fig 2). Lower assimilation rates observed in landraces under well-watered conditions, when compared with improved accessions, suggests a couple of possibilities. First, improved accessions likely undergo a more typical response to water stress, i.e. an overall reduction in photosynthetic activity. Second, landrace accessions have a lower photosynthetic capacity under ideal conditions but demonstrate more resilience to stress by maintaining similar rates of assimilation compared with their improved counterparts. This may be an example of differential plasticity between landraces and more improved accessions. For example, in other productivity traits, improved accessions are more productive under well-watered conditions, but experience a greater loss in response to water deficit than landraces, as seen in several species when compared to their wild counterpart [28]. It is also possible that level of domestication and breeding have contributed to morphological changes, which in-turn affect whole plant physiology. Examples of these morphological changes were reported by Milla and Matesanz [33], where domesticated plants, when compared with their wild counterparts, invested more resources into leaf production than stem production, while maintaining similar photosynthetic rates at the individual leaf. It may also be important to consider cultivation systems associated with landrace chile pepper. For example, landraces are frequently found in milpa polyculture systems, where increased branching could be favorable for improved light capture and microclimate management. However, additional study with emphasis on cultivation system is required to properly address this hypothesis. Finally, the relationship between precipitation parameters from the originating environment and primary branching variation in Mexican accessions further suggests that branching may be an adaptation associated with water availability (Table 5). Morphological adaptations associated with environment of origin have been identified in other studies, for example with stomatal size and density in Arabidopsis [32].

### Evidence of water deficit tolerant germplasm

We have identified traits in chile pepper that may provide adaptation to water deficit, such as branching and CO2 assimilation, which warrant further study. Some accessions expressed values of these traits that might make them relatively more drought tolerant or drought susceptible. For instance, the Mexican accession Ca0344 stands out as having exceptionally high productivity (measured as plant above-ground biomass), despite water deficit (75.7 g; Table 3) and maintains steady primary branching and CO_2_ assimilation rates when water deprived (Figs 1 and 2). On the other end of the spectrum, the paprika type, Szegedi 179, had very low biomass overall (49.6 g; Table 3) and had reduced branching and CO_2_ assimilation rates under water deficit (Figs 1 and 2). US accessions maintaining yield across irrigation treatments may be classified, for agronomic purposes, as water deficit tolerant (Fig 3). Most notable in this experiment was the New Mexico landrace, Canoncito, which has relatively high fruit yields and maintains fruit yield even when water deprived. This is considered a tolerant response, particularly when compared with a very susceptible accession, such as Anaheim M, which was the highest yielding accession under well-watered conditions, and experienced severe reduction in fruit weight under water deficit.

## Conclusion

With increasing risk of drought to agriculture, tolerance to soil water deficit is of the utmost importance. This study demonstrated differential response of above-ground traits to soil water deficit in diverse chile pepper germplasm selected from the U.S. and Mexico. Results provide opportunity for continued study in drought adaptation and soil water deficit tolerance in chile pepper, specifically through the identification of unique morphological and physiological responses to water deficit and high performing accessions.

## Acknowledgements

The authors would like to acknowledge members of the Mercer lab for assisting with experiment management and data collection. We would like to thank E. van der Knapp, A. Michel, J. Pérez-Alquicira, N. Taitano, R. Capouya, B. Pace, A. Aguilar Melendez, V. Bernau for assistance with chile pepper collections. We also appreciate feedback on earlier versions of this manuscript from N. Martínez, L. Connolly, and V. Bernau. We also than J. Vent for assistance in the greenhouse.

## Supporting information

**Figure 1**. Correlation matrix for seven environmental variables associated with origin of Mexican chile pepper landraces. Size and color indicate strength of the corelation (larger and darger blue is more positively correlated and larger and darger red is more negatively correlated). Variables, derived from BioClim and ISRIC, are: Total available soil water content, BIO12 = mean annual precipitation, BIO15 = precipitation seasonality, bIO16 = precipitation of the wettest quarter, BIO17 = precipitation of the driest quarter, BIO18 = precipitation of the warmest quarter, BIO19 = precipitation of the coldest quarter. Total available soil water content, BIO12, and BIO15 were maintained in the multiple regression model.

